# ERRγ deletion in podocytes accelerates aging related kidney disease

**DOI:** 10.64898/2026.05.11.724391

**Authors:** Xiaoxin Wang, Komuraiah Myakala, Natalia Shults, Rozhin Penjweini, Cheryl Clarkson-Paredes, Ewa Krawczyk, Sujit Hegde, Anastas Popratiloff, Julia Panov, Ruzong Fan, Garret Guthrie, Xiaoping Yang, Avi Rosenberg, Jay Knutson, Moshe Levi

## Abstract

We have recently demonstrated that treatment of aged mice with a pan-ERR agonist reverses age-related increase in urinary albumin, decrease in podocyte density, impaired mitochondrial function, and inflammation. The contribution of individual isoforms of ERRs however has not been determined. Since the aging kidney showed a possible compensatory increased expression of ERRγ in the podocytes, in the face of decreased ERRα expression, in the present study we aimed to determine the role of ERRγ in aging podocyte. To this end, we cross bred ERRγ floxed mice with podocin-Cre mice to achieve a podocyte-specific ERRγ deletion. While these mice at 3 months of age showed no effect on albuminuria compared to the wild type, when the mice were aged to 21 months of age, there was a significant increase in albuminuria and decrease in podocyte density. Furthermore, we found that the podocyte deletion of ERRγ primarily targeted the expression of mitochondrial biogenesis regulator PGC-1α, and mitochondrial fatty acid oxidation enzymes CPT1a and MCAD in the kidney. Electron Microscopy (EM) revealed thickened glomerular basement membrane and diffuse podocyte foot process effacement, as well as severe mitochondrial damage including cristae abnormalities, fragmentation, and changes indicative of altered fusion and fission dynamics. Fluorescence Lifetime Imaging Microscopy (FLIM) to determine NADH and FAD lifetimes indicate a metabolic shift from mitochondrial oxidative phosphorylation towards glycolysis, and decrease in mitochondrial redox capacity. Considering a significantly decreased expression of ERRα in aging podocytes plus its traditional role in mitochondrial function, these studies using podocyte ERRγ deletion suggested an overlapping mechanism for ERRα/ERRγ to act as modulators of age-related mitochondrial dysfunction and age-related kidney disease.

## INTRODUCTION

The fastest growing segment of the US population with impaired kidney function is the group aged ≥65 years. This population is expected to double in the next 20 years, whereas the number worldwide is expected to triple from 743 million in 2009 to 2 billion in 2050 (1). This will result in a marked increase in the elderly population with chronic kidney disease. This increase may be further amplified by other age-related comorbidities, including metabolic syndrome and hypertension, that accelerate age-related decline in renal function (2, 3).

A gradual age-related decline in renal function occurs even in healthy aging individuals (4–6) Progressive glomerular, vascular, and tubulointerstitial sclerosis is evident on renal tissue examination of healthy kidney donors with increasing age (7). In addition to aging, *per se*, metabolic syndrome and hypertension can induce mitochondrial dysfunction and inflammation, as well as endoplasmic reticulum stress, oxidative stress, altered lipid metabolism, and elevation of profibrotic growth factors in the kidney, which collectively contribute to age-related kidney disease (8–11).

There is variation in the rate of decline in renal function as a function of sex, race, and burden of comorbidities (12–17). Interestingly, examination of processes leading to renal sclerosis suggests a role for possible modifiable systemic metabolic and hormonal factors that can ameliorate the rate of sclerosis. In this regard, caloric restriction (CR) has been shown to decrease age-related kidney disease (18–20). We found that CR results in increased expression of the Estrogen Related Receptors (ERRs) in the kidney, therefore becoming a potential CR mimetic candidate (21).

The ERRs, ERRα (*NR3B1* and *ESSRA* genes), ERRβ (*NR3B2* and *ESRRB* genes), and ERRγ (*NR3B3* and *ESRRG* genes), are members of the nuclear receptor superfamily. ERRα and ERRγ are highly expressed in the kidney (22, 23).

In a recent study we determined that treatment of old mice with a pan-ERR agonist reversed age-related kidney disease, reversed the impairment in mitochondrial function, and reversed the increase in inflammation via the cGAS-STING pathway (21).

In that study we performed single-nuclei RNA sequencing to determine where *ERR*α and *ERR*γ mRNAs are expressed and regulated in the mouse kidney (21). We found that ERRα was expressed in most of the cell types within the kidney, most prominently in the proximal tubules, intercalated cells, and podocytes, and in many cases decreased with aging. Conversely ERRγ was detected in fewer cells and mainly in the proximal tubule and intercalated cells of young mice. Compared with young kidneys, the S1/S2 segments of aging proximal tubules showed a decline in both ERRα and ERRγ expression, whereas surprisingly ERRγ podocyte expression increased with aging (21). These results suggested that ERRγ may compensate for decreased ERRα in the podocytes.

The purpose of the present study was to determine the role of ERRγ in the glomerular podocytes of aging kidneys. We cross bred ERRγ floxed mice with podocyte Cre mice to achieve podocyte specific knockdown of ERRγ. We studied the mice when they were 21-month-old. The results indicate that ERRγ plays an important role in podocyte ultrastructure, mitochondrial integrity and function in mice as a function of aging.

## MATERIALS and METHODS

### Mice

All mice were maintained on a C57BL/6J background. ERRγ-floxed mice were generously provided by Dr. Anastasia Kralli at JHU. Podocyte-specific knockout was achieved by crossing floxed mice with NPHS2-Cre mice (Jax stock# 008205), which express Cre recombinase directed to podocytes. Experimental mice were housed and aged under a 12-h light/12-h dark cycle in a temperature-controlled environment, with ad libitum access to food and water. Animal studies and relative protocols were approved by the Animal Care and Use Committee at the Georgetown University. All animal experiment was conducted in accordance with the Guide for Care and Use of Laboratory Animals (National Institutes of Health, Bethesda, MD).

### Urine collection and analysis

Spot urine was collected before sacrifice and urine albumin and creatinine concentrations were assessed using kits from Exocell (Philadelphia, PA, USA) and BioAssay Systems (Hayward, CA, USA), respectively.

### RNA extraction and real-time quantitative PCR

Total RNA was isolated from the kidneys using Qiagen RNeasy mini kit (Valencia, CA), and cDNA was synthesized using reverse transcript reagents from Thermo Fisher Scientific (Waltham, MA). Quantitative real-time PCR was performed as previously described (24–27), and expression levels of target genes were normalized to 18S level.

### Immunohistochemistry

Formalin-fixed paraffin-embedded kidney sections were subjected to antigen retrieval with citrate buffer in high pressure heated water bath and staining was performed using anti-NPHS2 antibody (Catalog No. ab181143; Abcam, Waltham, MA) for 90 minutes; SignalStain^®^ Boost IHC Detection Reagent-HRP (Catalog No. 8114; Cell signaling technology, Danvers, MA) was applied followed by AEC chromogen. Imaging was done with MoticEasyScan Pro scanner (Motic, Vancouver, BC, Canada). Images were analyzed with HALO software (Indica Labs, Albuquerque, NM).

### Immunofluorescence microscopy

Frozen sections were used for immunostaining for synaptopodin (Catalog No. SE-19, Sigma, St. Louis, MO) and imaged with a laser scanning confocal microscope (SP8, Leica Biosystems, Buffalo Grove, IL). The expression level was quantified as sum of pixel values per glomerular area using ImageJ 1.44 software.

### FLIM setup for imaging of metabolic redox ratio

Two-photon FLIM was performed using an Olympus IX81/FV1000 confocal laser scanning microscope (Melville, NY, USA) equipped with a tunable Mai Tai BB DeepSee femtosecond laser (Spectra-Physics, Santa Clara, CA, USA) with wavelengths set to 860 and 750 nm for the excitation of FAD^+^ and NAD(P)H, respectively. The excitation light was passed through a 690 nm dichroic mirror and directed to an Olympus UPLXAPO 60×, 1.2 NA water immersion objective. An HPM-100-06 Hybrid detector (Becker & Hickl GmbH, Berlin, Germany) with bandpass filters 560/40 nm and 460/60 nm (Semrock BrightLine^®^, Rochester, NY, US) were used to detect FAD^+^ and NAD(P)H signals, respectively. Time-correlated single-photon counting (TCSPC) histograms were constructed using an SPC-180NX photon counting card (Becker & Hickl) and FLIM images were analyzed using SPC Image 8.8 software (Becker & Hickl GmbH).

### Electron Microscopy

Renal cortex tissues were fixed in the 2.5% glutaraldehyde/2%paraformaldehyde/ 0.05 mol/L cacodylate solution, post-fixed with 1% osmium tetroxide, and embedded in EmBed812. For *scanning electron microscopy (SEM)* evaluation, ultrathin sections (120 nm) were mounted in silicon wafers and observed with a Teneo LV FEG SEM (FEI, Thermo Fischer Scientific). For optimal results, we used the optiplan mode (high-resolution) equipped with an in-lens T1 detector (Segmented A+B, working distance of 8 mm). Low-magnification images (600×) were first taken for the observation then we generated high magnification images of our regions of interest (25,000) using 2 kV and 0.4 current landing voltage.

Glomerular basement membrane (GBM) thickness was measured using calibrated image analysis software (ImageJ). Scanning electron microscopy (SEM) images were acquired at ×25,000 magnification. The pixel-to-nanometer ratio was calibrated using the scale bar present on each image. GBM thickness was measured orthogonally to the capillary loop at multiple points per glomerulus, carefully avoiding oblique sections and regions with artifacts. For each capillary loop, 15 measurements were taken at regular intervals along the capillary wall. Measurements were collected from six glomerular capillary loops per glomerulus, resulting in 90 measurements per glomerulus (28). This sampling strategy provided a robust dataset for statistical analysis. To minimize the influence of occasional overestimations caused by oblique sections, the harmonic mean of all measured values was calculated for each glomerulus. This approach more accurately reflects the true membrane thickness across the capillary circumference, in accordance with stereological principles proposed by Weibel and Gomez (29). A total of ten glomeruli per animal and two animals per group were analyzed. All image analysis was performed under blinded conditions. Podocyte average foot process width (nm) measured by considering total number of foot processes/total length of the GBM (30). Podocyte foot process effacement (FPE) was quantified according to previous study reported in human glomerular disease (31).

The volume fraction of damaged mitochondria was determined using the morphometric technique with a dot grid. Mitochondrial damage was quantified by assessing abnormal mitochondria with irregular shape and size, signs of hypoplasia, inner and outer membrane disruption. Also, swollen cristae electron-lucent matrix, uneven thickening, homogenization, fragmentation together with cristae lose longitudinal orientation, tightness, and regular spacing. Filamentous mitochondria were determined as thin, highly elongated filament-like structures. Morphometric analysis was performed under blinded conditions by systematic uniform random sampling with the Fiji Software using 20 randomly selected images.

### Bulk RNA-seq

One microgram of total RNA samples was sent to Novogene (Sacramento, CA) for mRNA sequencing. RNA-seq fastQ files were filtered and trimmed from adaptors using Trimmomatic algorithm (32). The reads were aligned to *Mus musculus* genome assembly and annotation file GRCM38:mm10 using STAR algorithm (33). Gene expression was estimated in FPKM counts using RSEM algorithm (34). Differential expression was quantified with DeSeq2 algorithm (35). Absolute fold change of 1.5 and Bonferroni adjusted p-value of less than 0.05 was considered as significant change. All bioinformatics analysis was performed on T-BioInfo Platform (http://tauber-data2.haifa.ac.il:3000/). DAVID Bioinformatics (36, 37) and PANTHER Classification System (http://PANTHERdb.org/) were used to classify the differentially expressed genes into functional groups.

### Statistical analysis

Results are presented as the means ± SEM. Following the Grubbs’ outlier test, the data were analyzed by ANOVA and Newman–Keuls tests for multiple comparisons or by *t* test for unpaired data between two groups (Prism 6, GraphPad, San Diego, CA). Statistical significance was accepted at the *p*<0.05 level.

## RESULTS

### ERR**γ** deletion in podocytes worsens kidney injury in aged mice

We compared 21-month-old wild type mice (ERRγ-flox) and mice with podocyte-specific ERRγ deletion and found that podocyte-specific ERRγ deletion increased urinary albumin excretion, which is a marker of glomerular and podocyte injury (**Figure 1A**). The increase in albuminuria is likely related to worsened podocyte function as podocyte density represented by podocyte markers NPHS2 (podocin) **(Figure 1B)** and synaptopodin was further decreased in the kidneys with podocyte-specific ERRγ deletion (**Figure 1C**). Moreover, ultrastructural examination demonstrated increased glomerular basement membrane (GBM) thickening and podocyte foot processes effacement in aged kidneys which were worsened by the podocyte-specific ERRγ deletion (**Figure 2**). Taken together, we concluded that podocyte-specific ERRγ deletion accelerated the aging-related renal injury.

**Figure 1:**
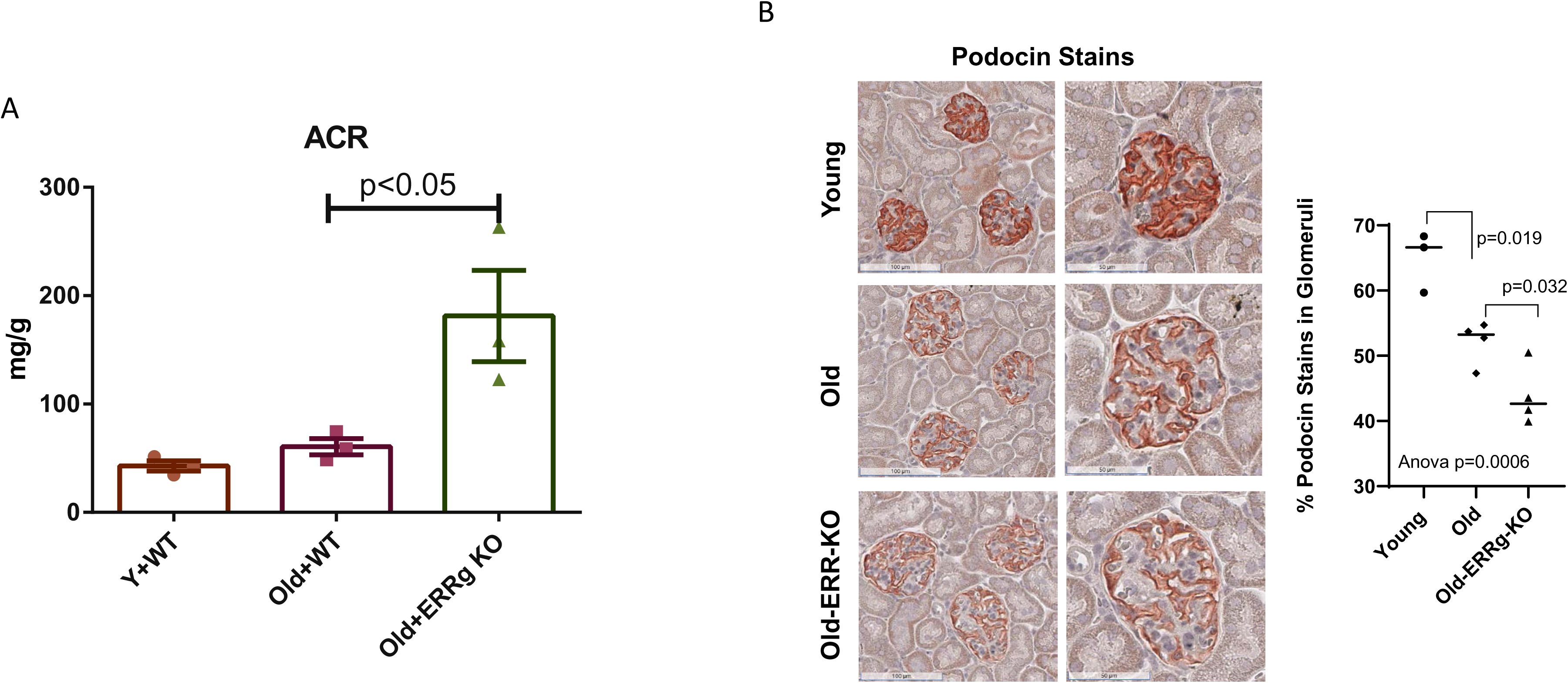
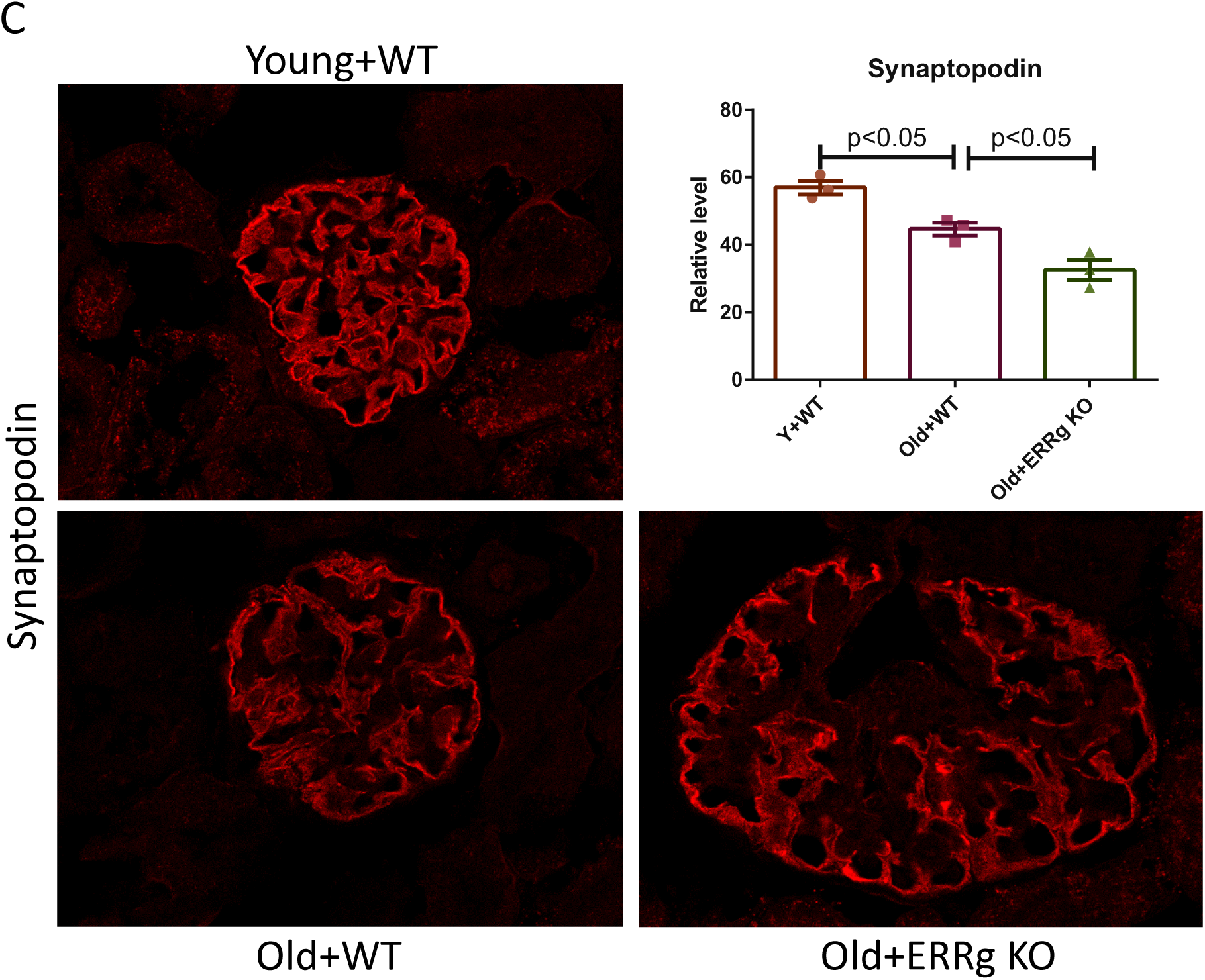
Podocyte-specific ERRγ knockout exacerbates albuminuria and decreases podocyte density in aged mice. **(A)** At 21 months of age, knockout mice exhibited significantly higher levels of albuminuria compared to age-matched wild-type controls (Old+WT). Both groups showed an increase in albuminuria relative to 3-month-old mice (Y+WT). (**B)** Immunohistochemistry of kidney sections for NPHS2 (podocin). **(C)** Immunofluorescence staining of kidney sections for synaptopodin. Data has been quantified using the ratio of total pixel intensity versus glomerular area. N=3-4 mice per group. Data were analyzed by ANOVA and Newman–Keuls tests.

**Figure 2:**
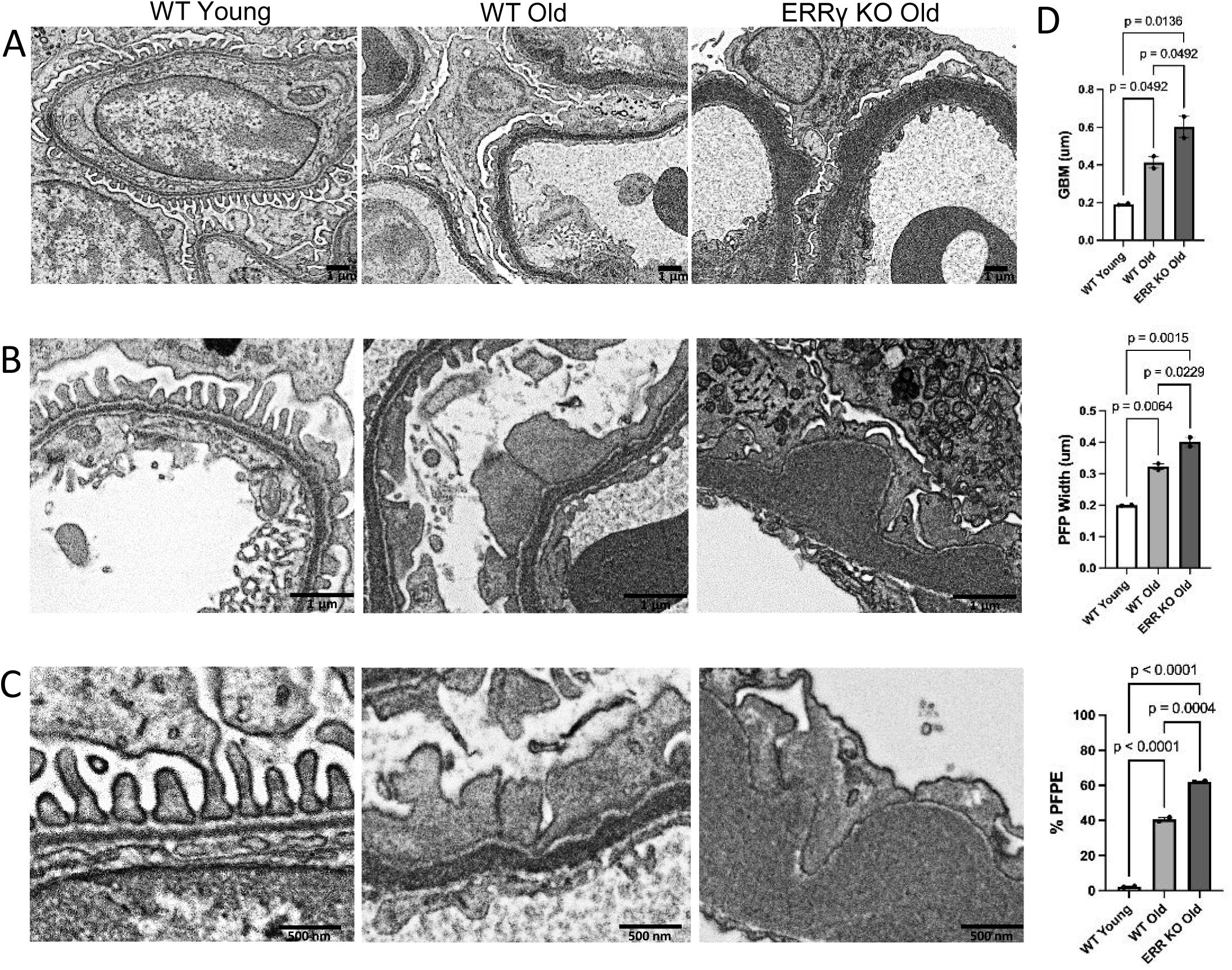
Ultrastructural changes in the glomerular filtration barrier. Electron microscopy reveals **(A)** normal thickness glomerular basement membrane (GBM) in young wild-type (WT) mice. In aged WT mice, the GBM is thickness is increased compared with young WT animals. In aged ERRγ deletion (KO) mice, the GBM is significantly thicker than in aged WT mice. Magnification ×25,000. **(B)** Young WT mice exhibit regularly spaced and distinct podocyte foot processes. In contrast, aged WT mice demonstrate widening and fusion of PFPs. In aged ERRγ KO mice, these changes are exacerbated, with pronounced podocyte foot process effacement (PFPE). Magnification ×65,000. **(C)** Higher magnification shows: In WT mice, podocyte foot processes are connected by intact slit diaphragms (arrow). In aged WT mice, slit diaphragms are partially lost, with a reduced frequency of visible filtration pores compared to young WT animals. This alteration is aggravated in aged ERRγ KO mice. Magnification ×80,000. **(D)** Quantitative analyses demonstrate glomerular structural alterations in young WT, aged WT, and aged ERRγ KO groups. N= 3 mice per group. Data were analyzed by ANOVA and Newman–Keuls tests.

### ERR**γ** deletion in podocytes worsens the mitochondrial morphology and function

Aging kidneys manifest mitochondrial dysfunction through impaired energy metabolism, altered morphology, and suppressed oxidative phosphorylation (OXPHOS). To quantify these shifts, we employed label-free fluorescence lifetime imaging microscopy (FLIM) to determine the ratio of free to bound NAD(P)H, providing a robust metabolic estimate of glycolysis versus OXPHOS. A lower free-to-bound NAD(P)H ratio serves as a marker for enhanced oxidative phosphorylation and reduced glycolysis. Conversely, an elevated free-to-bound NAD(P)H ratio signifies a metabolic shift toward glycolysis and diminished mitochondrial efficiency (38). We resolved these shifts by leveraging the distinct autofluorescence decay rates of NAD(P)H and the relative intensities of both cofactors. We observed a significant increase in the free-to-bound NADH ratio in kidney sections from aged podocyte-specific ERRγ deletion mice compared to aged wild type ones. However, at 21 months of age, the wild type aged mice had no significant difference from young mice (**Figure 3**).

**Figure 3:**
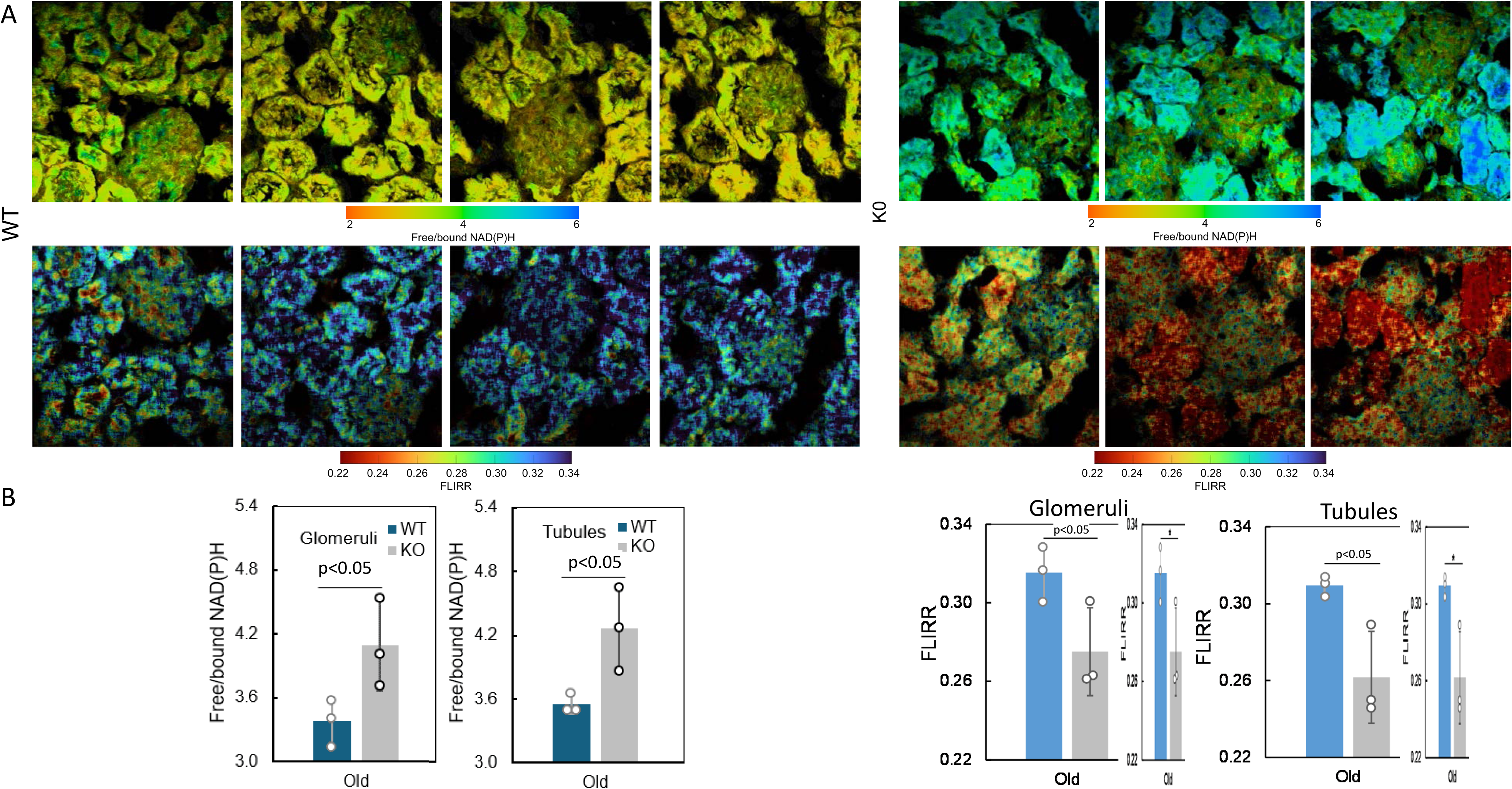
FLIM imaging of kidneys from podocyte-specific ERRγ knockout and wild-type mice. **(A)** Pseudocolor mapping of free/bound NAD(P)H and FLIRR in the interstitial environment of aged podocyte-specific ERRγ deletion mice and aged wild type mice. In the color, red indicates lower values, whereas blue indicates higher values. **(B)** Bar charts of the free/bound NAD(P)H and FLIRR; error bars are the standard deviation and individual data points are technical/biological replicates. N=3 mice per group. Data were analyzed by ANOVA and Newman–Keuls tests.

Changes in the metabolic redox ratio can be attributed to a change in the free/bound NAD(P)H ratio but is also evidenced by a change in the relative amount of NAD(P)H/NAD^+^ or fluorescence lifetime based redox ratio (FLIRR); FLIRR is calculated from the bound decay components of NAD(P)H and FAD^+^ (39). FLIRR is independent of the concentration, laser power, focus, and many other instrument parameters and is thus considered a robust indicator of the metabolic state (39). Based on the results shown in **Figure 3**, a consistent lower FLIRR was measured for kidney sections from aged podocyte-specific ERRγ deletion mice when compared to aged wild type ones. A reduced FLIRR reflects dominance of glycolysis in aged podocyte-specific ERRγ deletion mice, which is in agreement with free/bound NAD(P)H results.

Next, we used scanning electron microscopy to determine mitochondria morphology. In podocytes from young mice, mitochondria showed organized cristae in uniformly electron-dense matrix. Thin, filament-like mitochondria were observed in young WT mice, consistent with normal fusion–fission activity. In aged podocytes, mitochondria exhibited cristae disruption, including fragmentation and homogenization, along with a modest increase in filament-like mitochondria. In aged podocytes with ERRγ deletion, mitochondrial damage is more pronounced and showed a significant increase in thin, filament-like mitochondria accumulating within damaged mitochondria (**Figure 4**).

**Figure 4:**
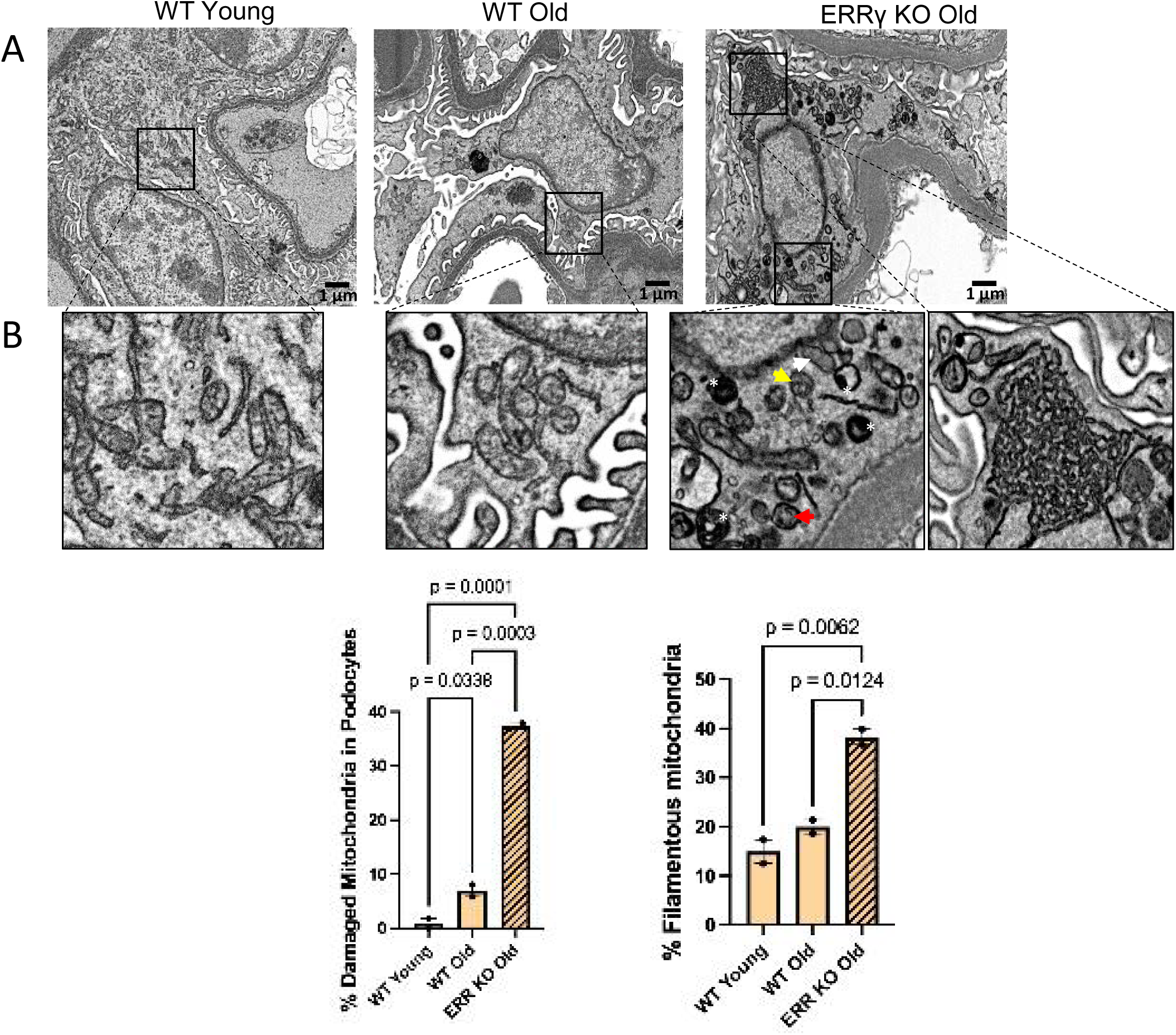
Mitochondrial ultrastructural changes in podocytes with aging and ERRγ deficiency. **(A)** Electron microscopy images demonstrate mitochondria distribution and organization in young WT and KO and in aged ERRγ KO mice. Magnification x25,000. **(B)** Under higher magnification in young WT mice mitochondria in oval or elongated shape with organized cristae in electron-dense matrix. Thin mitochondria, filament-like structures were also observed (arrows). In aged WT mice, mitochondria exhibit cristae disruption: fragmentation and homogenization (arrows), along with a modest increase in filament-like mitochondria compared with young WT. In aged ERRγ KO mice, mitochondrial damage is pronounced. Arrows indicate cristae abnormalities: (1) crystallization (red), (2) fragmentation (yellow), and (3) homogenization (white). Asterisks mark degraded mitochondria within autophagosomes, indicating increased mitophagy. Notably, ERRγ KO podocytes show a significant increase in filament-like mitochondria accumulating within damaged mitochondria. N=3 mice per group. Data were analyzed by ANOVA and Newman–Keuls tests.

Consistent with the above morphology observations, we found the expression level of PGC-1α, which mediates mitochondrial biogenesis, and CPT1a, and MCAD, which mediate mitochondrial fatty acid oxidation, was decreased more in aged podocyte-specific ERRγ deletion kidneys than the wild type aged kidneys (**Figure 5**).

**Figure 5:**
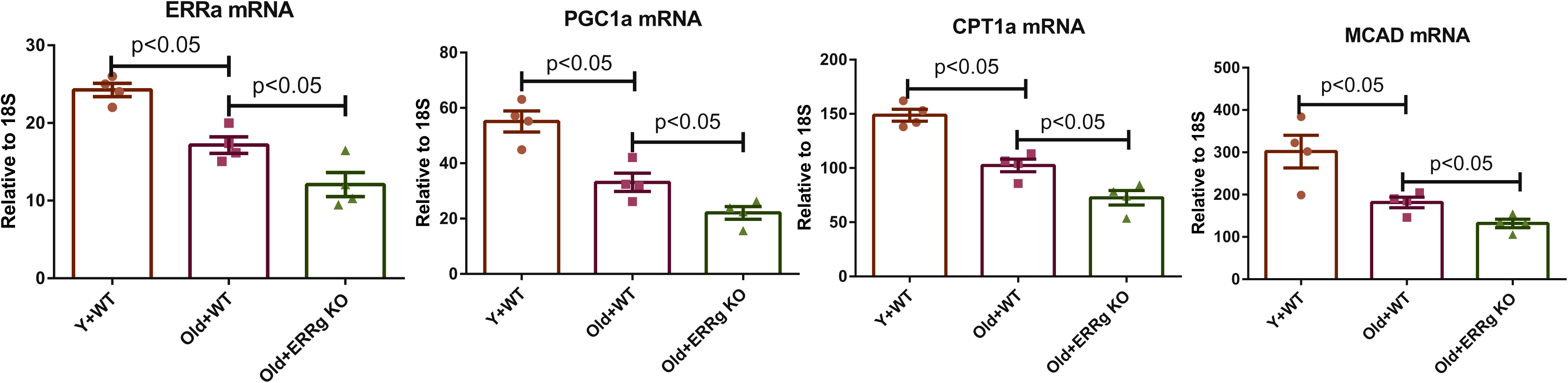
Podocyte-specific ERRγ deletion exacerbates the age-related decline of mitochondrial metabolic mediators. Quantitative PCR analysis of mRNA expression for *Err*α*, Pg1*α*, Cpt1a, and Mcad* in the kidneys of young and aged mice. These markers were significantly downregulated in aged wild-type kidneys compared to young controls, and further suppressed in aged ERRγ knockout mice. N=4 mice per group. Data were analyzed by ANOVA and Newman–Keuls tests.

### ERR**γ** podocyte deficiency orchestrates a metabolic-to-mesenchymal transition in aging kidneys

To evaluate the transcriptomic impact of ERRγ loss in the aging podocytes, we performed RNA-seq analysis comparing aging wild type kidneys and aging kidneys with podocyte-specific ERRγ deletion (KO). Gene set enrichment analysis (GSEA) revealed a profound collapse of the mitochondrial transcriptional program in KO kidneys, characterized by the significant downregulation of modules associated with oxidative phosphorylation (OXPHOS), the electron transport chain, and mitochondrial inner membrane integrity. This mitochondrial decay was accompanied by a sharp upregulation of genes involved in cell migration and extracellular matrix organization, alongside signatures of epithelial-to-mesenchymal transition (EMT). Notably, the upregulation of the “Rodwell aging kidney” gene set suggests that the loss of ERRγ significantly accelerates the molecular clock of renal senescence, driving podocytes away from their specialized epithelial identity toward a migratory, dysfunctional phenotype (40–43) (**Figure 6**).

**Figure 6.**
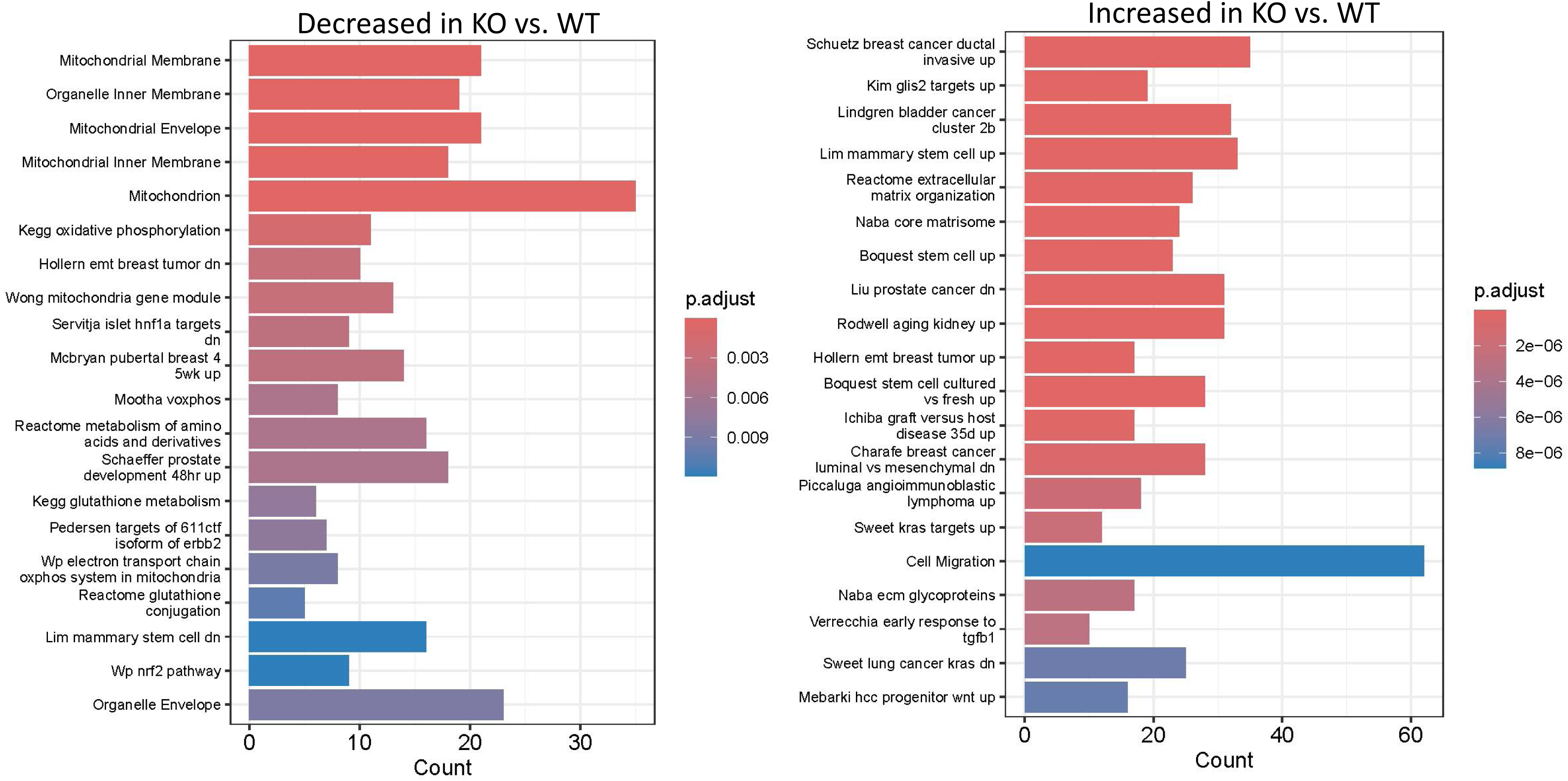
RNA-seq pathway enrichment analysis in aged podocyte-specific ERRγ knockout kidneys. Bar plots showing significantly enriched pathways and gene sets derived from differential expression analysis comparing aged podocyte-specific ERRγ knockout kidneys with aged wild-type controls. The left panel shows pathways significantly enriched among downregulated genes in ERRγ-deficient kidneys. The right panel shows pathways enriched among upregulated genes. Bar length represents the number of overlapping genes within each pathway or gene set, and color indicates the adjusted p value.

## DISCUSSION

Estrogen-related receptor gamma (ESRRG, ERRγ) has emerged as a central transcriptional regulator of mitochondrial metabolism across multiple high-energy tissues. In the kidney, earlier work by Zhao et al. (44) demonstrated that ERRγ is highly expressed in renal epithelial cells and is essential for coordinating mitochondrial oxidative phosphorylation (OXPHOS) and renal tubular reabsorptive functions through both HNF1β-dependent and independent transcriptional programs. However, that study also revealed that ERRγ expression is largely restricted to tubular compartments, with minimal expression detected in glomerular cells, leaving its role in podocytes unresolved.

In the present study, we provide the first in vivo evidence that ERRγ is essential for maintaining podocyte integrity, particularly in the context of aging. Using a podocyte-specific ERRγ deletion model, we demonstrate that loss of ERRγ accelerates age-associated kidney injury, characterized by increased albuminuria, glomerular basement membrane thickening, and podocyte foot process effacement. These findings extend the role of ERRγ beyond tubular compartments and identify it as a critical regulator of glomerular function in aging kidney.

Our findings align with and extend previous work from our group demonstrating that ERR signaling is a central modulator of mitochondrial dysfunction and inflammation in the aging kidney (21). In that study, we showed that both ERRα and ERRγ expression were reduced in aging human and mouse kidney tubules, and that pharmacological activation of ERRs using a pan-ERR agonist reversed key features of kidney aging, including albuminuria, podocyte loss, mitochondrial dysfunction, and inflammatory signaling. Notably, ERR agonism restored mitochondrial gene expression, improved mitochondrial oxygen consumption rate (OCR), and suppressed cGAS-STING mediated inflammation, highlighting ERR signaling as a critical link between mitochondrial health and inflammaging.

Importantly, the current study provides mechanistic insight into the cell-type specific role of ERRγ within the glomerular podocyte cells. While our previous work demonstrated that ERR agonism improves podocyte density and kidney function at the whole-organ level, it remained unclear whether these effects were mediated directly through the podocytes or indirectly via tubular or systemic mechanisms. By selectively deleting ERRγ in podocytes, we now demonstrate that ERRγ is intrinsically required for maintaining podocyte metabolic function and structural integrity.

A major finding of this study is that ERRγ deficiency induces profound metabolic reprogramming in podocytes. Earlier study by Zhao et al showed that mitochondrial dysfunction precedes structural and functional kidney abnormalities in tubular ERRγ-deficient mice (44). Interestingly, our data reveal a similar paradigm in podocytes. Using florescence life time imaging microscopy (FLIM), we show that loss of ERRγ shifts podocyte metabolism from oxidative phosphorylation toward glycolysis, consistent with impaired mitochondrial efficiency. This is accompanied by marked mitochondrial structural abnormalities and downregulation of key regulators of mitochondrial biogenesis and fatty acid oxidation, including PGC-1α, CPT1a, and MCAD. These findings are highly consistent with our previous observations that ERR signaling regulates mitochondrial gene networks in the aging kidney and extend this concept to a podocyte-specific context.

Although ERRα and ERRγ share overlapping transcriptional targets and are often considered functionally redundant, our data suggest that ERRα is unable to compensate for the loss of ERRγ in podocytes. One important explanation is that ERRα expression is already reduced in podocytes under aging, as reported in our prior study (21), thereby limiting its compensatory capacity. In contrast, ERRγ appears to be relatively preserved or even upregulated in podocytes, potentially serving as the dominant ERR isoform maintaining mitochondrial gene expression in this cell type. Under these conditions, ERRγ may function as the primary metabolic regulator, while ERRα plays a more minor or permissive role. Consequently, deletion of ERRγ removes the predominant transcriptional driver of mitochondrial function, and the already diminished ERRα pool is insufficient to sustain metabolic homeostasis. As a result, mitochondrial oxidative phosphorylation is impaired, forcing podocytes to rely more heavily on glycolysis as an energy source. However, glycolysis is insufficient to meet the high energy demands of podocytes, leading to an overall energy deficit, cellular dysfunction, and eventual structural damage.

In addition to metabolic dysfunction, our transcriptomic analyses reveal that ERRγ loss drives a transition toward a pro-fibrotic and dedifferentiated state, with upregulation of extracellular matrix organization, cell migration, and epithelial-to-mesenchymal transition (EMT)-related pathways. Notably, enrichment of aging-associated gene signatures suggests that ERRγ deletion accelerates molecular aging processes within podocytes. These findings parallel our prior observations that mitochondrial dysfunction in aging kidneys is tightly coupled to inflammation and senescence pathways, further supporting a model in which ERRγ acts as a key regulator of the metabolic-inflammatory axis in renal aging (21).

Taken together, our findings support a model in which ERRγ serves as a dominant regulator of metabolic homeostasis in podocytes in an age-dependent manner. In young mice, ERRγ expression is relatively low compared to ERRα, suggesting that ERRα may play a more prominent role under basal conditions. However, during aging, ERRα expression declines, while ERRγ becomes relatively more important for maintaining mitochondrial function. This shift creates a context in which podocytes become increasingly reliant on ERRγ, with limited compensatory capacity from ERRα. Consequently, loss of ERRγ has minimal impact in young animals but leads to pronounced metabolic insufficiency, activation of fibrotic and inflammatory pathways, and accelerated glomerular aging in aged mice.

Overall, this study establishes a previously unrecognized, cell type–specific role for ERRγ in maintaining podocyte metabolic homeostasis and provides new mechanistic insight into how mitochondrial regulation is altered during aging. Our findings define how ERRγ functions within the broader ERR network to sustain podocyte integrity under age-associated stress. We demonstrate that the requirement for ERRγ is highly context-dependent, emerging specifically in aging when compensatory capacity from ERRα is diminished. This work therefore uncovers a key mechanism by which transcriptional control of mitochondrial metabolism becomes vulnerable in podocytes, linking ERRγ-dependent metabolic regulation to glomerular aging and injury. These insights refine our understanding of ERR biology in the kidney and provide a conceptual framework for how metabolic regulators drive cell-type–specific susceptibility in chronic kidney disease.

## Acknowledgements

The studies in this manuscript were supported by an NIH R01 DK127830 grant to ML.

